# Deconvoluting fiber type proportions from human skeletal muscle transcriptomics and proteomics data using FibeRtypeR

**DOI:** 10.1101/2025.09.17.676778

**Authors:** Thibaux Van der Stede, Roger Moreno-Justicia, Freek Van de Casteele, Eline Lievens, Alexia Van de Loock, Jonas Vandecauter, Peter Merseburger, Jimmy Van den Eynden, Delphi Van Haver, An Staes, Simon Devos, Morten Hostrup, Pieter Mestdagh, Jo Vandesompele, Ben Stocks, Atul S Deshmukh, Wim Derave

**Affiliations:** Department of Movement and Sports Sciences, Ghent University, Ghent, Belgium; Unit for Molecular Immunology and Inflammation, VIB-UGent Center for Inflammation Research, Flanders Institute for Biotechnology (VIB), Ghent, Belgium; Department of Internal Medicine and Pediatrics, Ghent University, Ghent, Belgium; Novo Nordisk Foundation Center for Basic Metabolic Research, Faculty of Health and Medical Sciences, University of Copenhagen, Copenhagen, Denmark; Department of Human Structure and Repair, Ghent University, Ghent, Belgium; Cancer Research Institute Ghent (CRIG), Ghent, Belgium; VIB Proteomics Core, Ghent, Belgium; Department of Biomolecular Medicine, Ghent University, Ghent, Belgium; Department of Nutrition, Exercise and Sports, University of Copenhagen, Copenhagen, Denmark; Department of Molecular Medicine and Surgery, Integrative Physiology, Karolinska Institutet, Stockholm, Sweden

**Keywords:** Skeletal muscle, fiber type, transcriptomics, proteomics, deconvolution, FibeRtypeR

## Abstract

Muscle fibers are the dominant, multinucleated cell type in skeletal muscle. In humans, they can be classified as slow (type 1) and fast (type 2) fibers, traditionally based on distinct properties of their contractile machinery. Slow and fast fibers are characterized by shared and specific cellular complexities that are pivotal for their adaptive capacity to exercise or role in metabolic disease progression. In this era of omics research, there is a critical need to accurately infer muscle fiber type proportions from bulk tissue omics datasets, as single-fiber approaches are not feasible in large-scale or retrospective studies. Here we present FibeRtypeR, an easy-to-use web application to accurately estimate fiber type proportions from bulk transcriptomics and proteomics datasets: *https://muscleapps.ugent.be*. FibeRtypeR exploits transcriptomics and proteomics profiles of 1000 fiber-typed individual human skeletal muscle fibers as a reference dataset and is validated against paired immunohistochemical fiber type determinations in 160 muscle biopsies. We show that FibeRtypeR can be applied to public datasets, illustrating the application potential across a wide range of biological contexts such as aging, disease and exercise training. This new freely accessible computational tool will prove valuable to the skeletal muscle research community.

**Key points:**

- Bulk muscle omics datasets lack fiber type specific information
- Our new tool, FibeRtypeR, leverages in-house collected single-fiber profiles allowing for accurate fiber type inference
- FibeRtypeR is methodologically robust across omics technologies and workflows
- We host FibeRtypeR as an intuitive open-access Shiny app, applicable to new and publicly available transcriptomics and proteomics datasets

## Introduction

As a heterogeneous tissue, skeletal muscle is a complex amalgamation of cell types, each with a unique role within skeletal muscle structure and function. The muscle fibers, long multinucleated cells that comprise up to 90% of the total muscle volume, play a pivotal role in locomotion via the contractile muscle machinery. Myosin heavy chain (MYH) is a central motor protein for muscle contraction and expression of specific MYH isoforms (MYH1, MYH2 and MYH7) is traditionally used to classify fast and slow fibers (Billeter et al., 1981; Schiaffino and Reggiani, 2011). Nonetheless, the differences between fiber types are broader than their contractile protein expression, as slow fibers display a higher resistance to fatigue, exhibit a higher mitochondrial content and rely more on their oxidative metabolism than their fast counterparts (Bottinelli et al., 1996; Garnett et al., 1979; Saltin and Gollnick, 1983; Zierath and Hawley, 2004).

The proportion of different fiber types in human skeletal muscle shows great inter-individual variation and is to a large extent genetically determined (Simoneau and Bouchard, 1995). A lower proportion of slow fibers has been associated with metabolic and functional abnormalities, such as increased risk for insulin resistance and obesity (Blackwood et al., 2022; Serrano et al., 2023). Fiber type distribution is also relevant for sports performance, as it could be used for talent identification in different sports (Saltin et al., 1977) and individualization of training and recovery strategies (Bellinger et al., 2020; Lievens et al., 2020; Trappe et al., 2000; Van de Casteele et al., 2024a). In recent years, various omics technologies have been on the rise aiming to expand our understanding of the complex molecular regulation of muscle adaptations to exercise (Blazev et al., 2022; Hoffman, 2017; Hostrup et al., 2022; Pillon et al., 2020), whereby only few studies investigated fiber type differences (Deshmukh et al., 2021; Moesgaard et al., 2024; Reisman et al., 2024; Van der Stede et al., 2024; Wyckelsma et al., 2025).

This is in part due to most current omics approaches being performed on bulk muscle, which is a homogenate of all residing cell types and makes it difficult to estimate the precise fiber type composition. Fiber type proportions in omics datasets are relevant and should be included in analysis that compare between experimental groups or for statistical modelling (e.g., as covariate) of the large variation between biopsies within the same individual (Horwath et al., 2021). The two gold standard methodologies that are commonly applied to tackle these questions are histochemical staining and antibody-based identification of fiber types within muscle cross sections or single-fiber SDS-PAGE analysis after manual isolation of individual muscle fibers from fresh or freeze-dried muscle biopsies (Bloemberg and Quadrilatero, 2012; Murach et al., 2019). These methods allow for accurate fiber typing, but are time-consuming and require additional tissue material for cryosectioning or fiber isolation. Furthermore, they do not provide exact quantitative information of the marker expression. On top, often only a single marker, myosin heavy chain isoform, is considered when inferring muscle fiber type, which can lead to misinterpretation of the true muscle fiber type heterogeneity (Moreno-Justicia et al., 2025).

The development of computational approaches to accurately quantify fiber type proportions from bulk omics datasets would therefore be a major step forward. This concept, termed computational cellular deconvolution, has been an important focus in the bioinformatics community with regular introduction of new tools (Avila Cobos et al., 2020). Recent iterations of these tools exploit the power of single-cell RNA sequencing as these use the cell type specific transcriptional profiles as reference input in the deconvolution algorithm (Avila Cobos et al., 2023; Tsoucas et al., 2019; Wang et al., 2019). This concept is yet to be transferred to the skeletal muscle field, as standard single-cell omics workflows are unable to measure the large and multinucleated skeletal muscle fibers. Indeed, a recent endeavour to use cell type deconvolution in the context of fiber typing utilized single-nuclei RNA sequencing data as reference (Oskolkov et al., 2022), which given the multinucleated nature of muscle fibers, might not be the most suitable approach. Moreover, this tool is limited to only transcriptomics analysis.

Recent technological advances in the fields of transcriptomics and mass spectrometry-based proteomics have enabled the unbiased measurement of isolated muscle fibers, being able to fiber type and explore the true complexity of their biological processes at the same time (Murgia et al., 2021). In fact, we and others have exploited this technological potential to uncover muscle fiber heterogeneity and fiber type-specific effects of exercise training, aging, or nemaline myopathy (Hostrup and Deshmukh, 2025; Moreno-Justicia et al., 2025; Murgia et al., 2017). Here, we present FibeRtypeR, a fiber type deconvolution tool that can process skeletal muscle bulk transcriptomics or proteomics datasets as an input and provide an accurate estimation of the fiber type ratio of the provided samples. FibeRtypeR can process transcriptomics and proteomics data, as the tool uses two distinct reference datasets that include data from 1000 single muscle fibers from multiple donors (Moreno-Justicia et al., 2025). In addition, FibeRtypeR is available as an open-source, user-friendly shiny app (https://muscleapps.ugent.be). FibeRtypeR facilitates the investigation of muscle fiber type composition across a range of biological and clinical settings, particularly when sample availability is limited. Moreover, the tool can retrospectively be applied to published datasets, where fiber type could not be evaluated previously.

## Results

### FibeRtypeR accurately infers fiber type proportions

A schematic overview of the FibeRtypeR deconvolution approach is displayed in **Figure 1**. FibeRtypeR was developed based on transcriptome and proteome profiles from over 1000 single muscle fibers isolated from *vastus lateralis* biopsies collected from multiple donors (n=12M/2F and n=5M for transcriptomics and proteomics, respectively) (**Figure 1A**) (Moreno-Justicia et al., 2025). Based on their transcriptome- and proteome-wide characteristics, these fibers separated into two distinct clusters, corresponding to slow and fast muscle fibers (Moreno-Justicia et al., 2025). Using this reference atlas, FibeRtypeR estimates fiber type proportions in new experimental datasets by comparing bulk transcriptomics or proteomics profiles to the fiber-type-specific genes and proteins (**Figure 1B**). To validate its accuracy, FibeRtypeR was benchmarked by comparing fiber types deconvoluted from bulk muscle transcriptomics and proteomics data generated from *vastus lateralis* and *gastrocnemius* biopsies of young, healthy and recreationally active individuals (n=20M/20F) (**Figure 1B**) with paired immunohistochemistry (IHC)-based fiber typing (using myosin isoforms) on cryosections of the same muscle biopsies (**Figure 1C**). The FibeRtypeR prediction showed strong significant correlations between IHC and both the transcriptomics and proteomics data in the *vastus lateralis* (r = 0.82 and 0.86; p < 0.001) and *gastrocnemius* (r = 0.81 and 0.82; p < 0.001) muscle (**Figure 2A**). Similarly, the transcriptome- and proteome-based estimations showed strong positive correlations with each other (r = 0.77 – 0.79; p < 0.001) (**Figure 2B**). Bland-Altman analysis further confirmed strong agreement between FibeRtypeR and IHC proportions for slow (or fast) fibers for the transcriptomics workflow, whereas a greater proportion of slow fibers (by approximately 9-10 percentage points) was determined by the proteomics workflow (**Figure 2C**).

**Figure 1.**
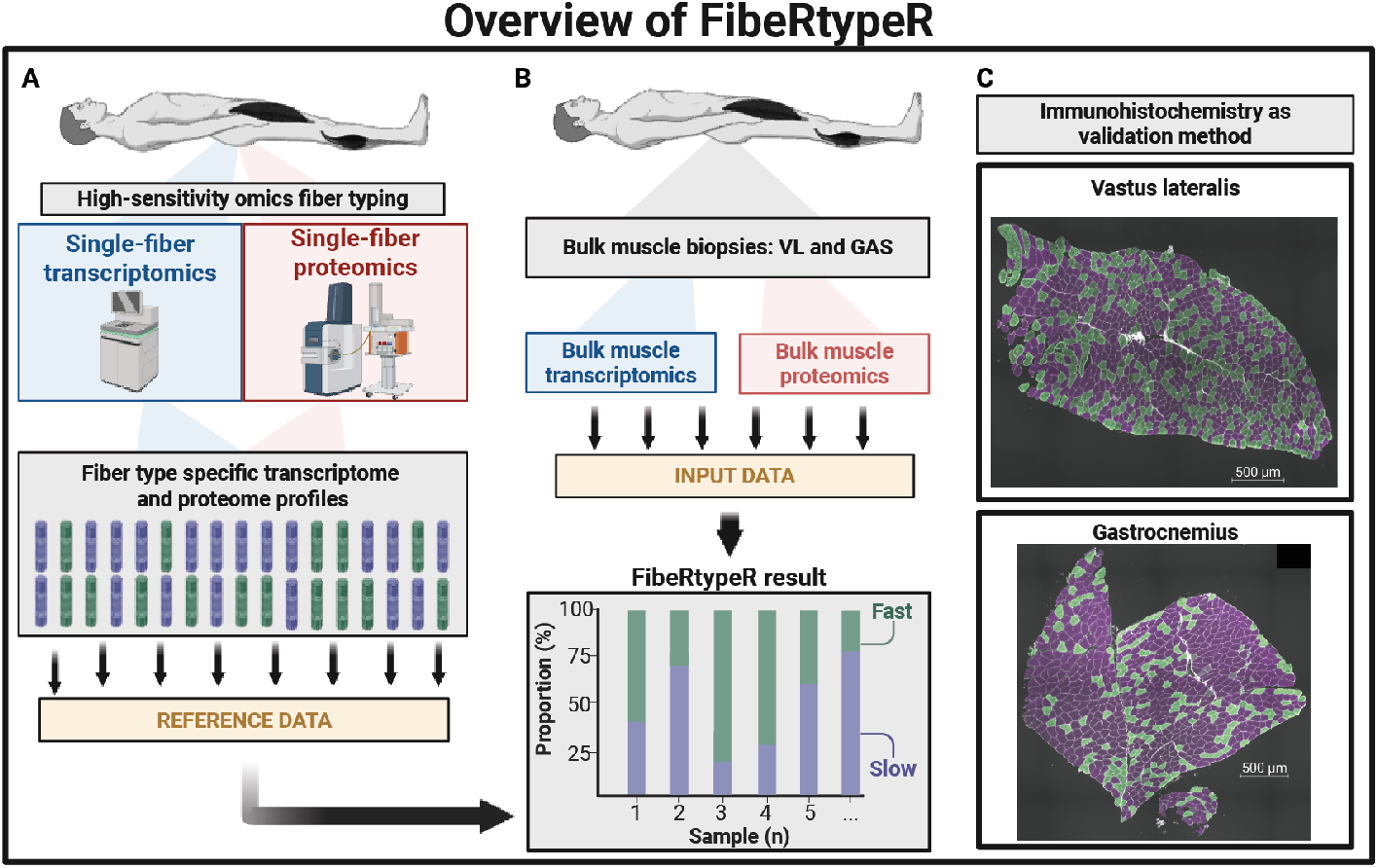
Graphical overview of FibeRtypeR. (**A**) Single-fiber transcriptome and proteome profiles are used as reference dataset for the deconvolution algorithm. (**B**) Collection of muscle biopsies and subsequent bulk transcriptome and proteome analysis. FibeRtypeR then infers fiber type proportions on this input dataset. VL: vastus laterals; GAS: gastrocnemius. (**C**) Computational inference of fiber type proportions from bulk muscle were compared to IHC-based determinations of fiber type (based on MYH isoforms). Representative images of immunohistochemical evaluation of fiber type in muscle cross-sections are shown.

**Figure 2.**
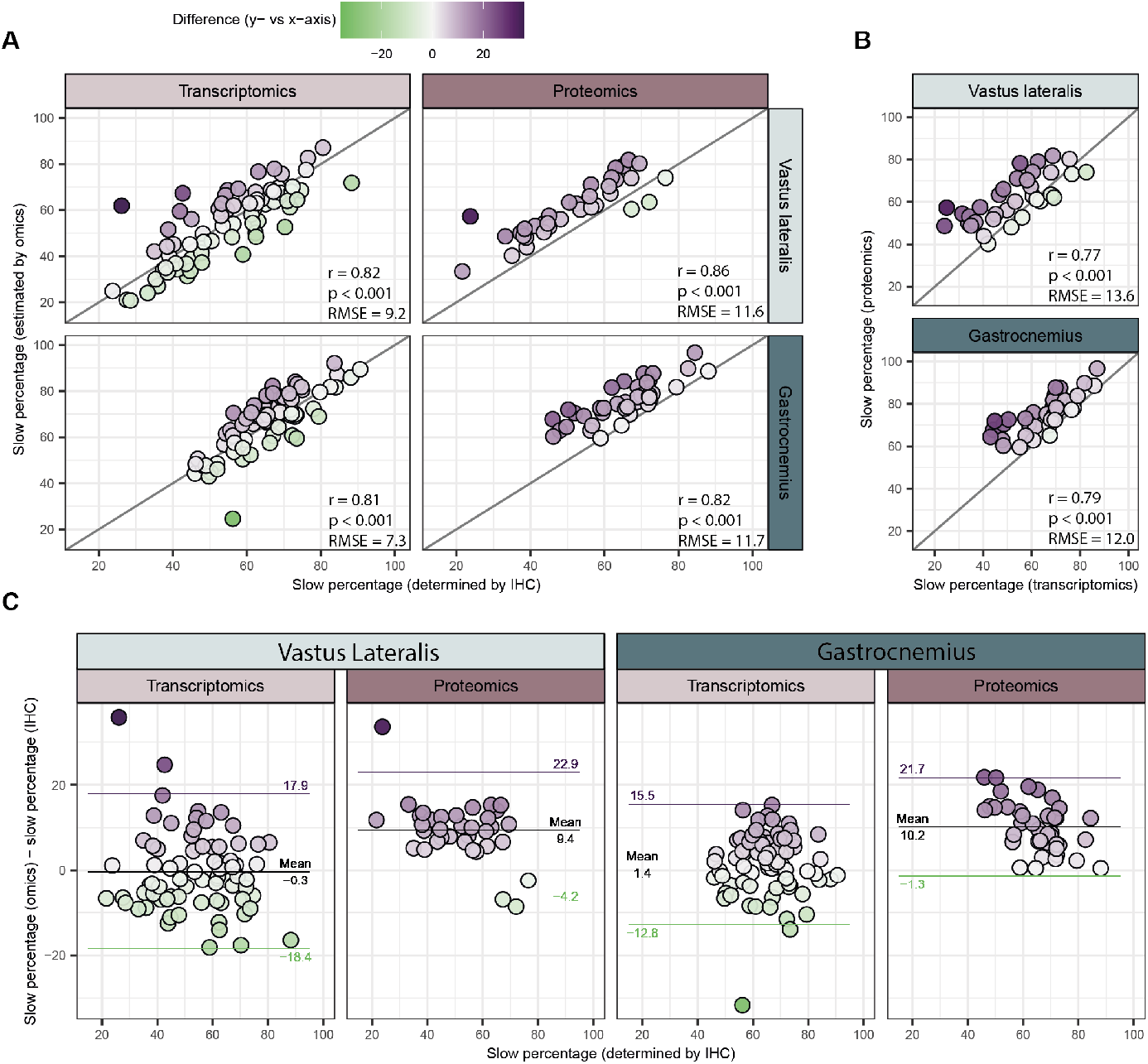
Validation of FibeRtypeR. (**A**) Correlation between fiber type proportions from IHC (x-axis) and transcriptomics and proteomics technologies (y-axis-, separately for vastus lateralis and gastrocnemius samples). Diagonal lines are lines of identity. (**B**) Correlation between transcriptomics and proteomics, separately for vastus lateralis and gastrocnemius samples. Diagonal lines are lines of identity. (**C**) Bland-Altman plots illustrating accuracy of fiber type inferences. Purple and green lines are 95% limits of agreement.

### FibeRtypeR draws from a set of genes and proteins beyond myosin heavy chains

FibeRtypeR infers muscle fiber type proportions by integrating multiple genes and proteins, selected based on log_2_-fold changes (LFC) of these features between individual slow (type 1) and fast (type 2A, 2A/X, 2X) *vastus lateralis* fibers. To benchmark the optimal threshold for feature selection, we systematically tested different LFC cut-offs (1, 2, 3 or 4) and evaluated performance based on correlation coefficients and root mean square error (RMSE) between deconvoluted and IHC-based fiber type estimates (**Figure 3A**). For the transcriptomics dataset, correlation coefficients were overall similar for the *vastus lateralis* muscle, although the lowest RMSE values were achieved with a LFC of 1 for both *vastus lateralis* and *gastrocnemius* muscle. For proteomics data, correlation coefficients did not differ substantially between the different thresholds, but the best RMSE value was achieved with a LFC cut-off of 2 (**Figure 3A**). These LFC thresholds resulted in the selection of 65 and 18 marker genes and proteins for slow fibers and 52 and 18 marker genes and proteins for fast fibers in transcriptomics and proteomics, respectively (**Figure 3B and Table S1**). To improve accuracy, we excluded genes (n = 13, e.g., *IGFN1, IRX3, PFKFB2* and *XIRP1*) that we previously identified to be responsive to acute exercise in bulk muscle (Van der Stede et al., 2025). There was only a modest overlap of selected features between the transcriptomics and proteomics datasets, indicating the importance of specificity for marker selection at the RNA or protein level (**Figure 3C**). Of note, the selected features belong to distinct biological categories, including contraction (e.g., MYH2, TNNT3 and MYL3), metabolism (e.g., LDHB and *CKB*), ubiquitin regulation (e.g., USP28 and USP48) and non-coding RNAs (e.g., *LINC01405*). This functional diversity highlights the capacity of FibeRtypeR to capture muscle fiber heterogeneity beyond traditional contractile elements.

**Figure 3.**
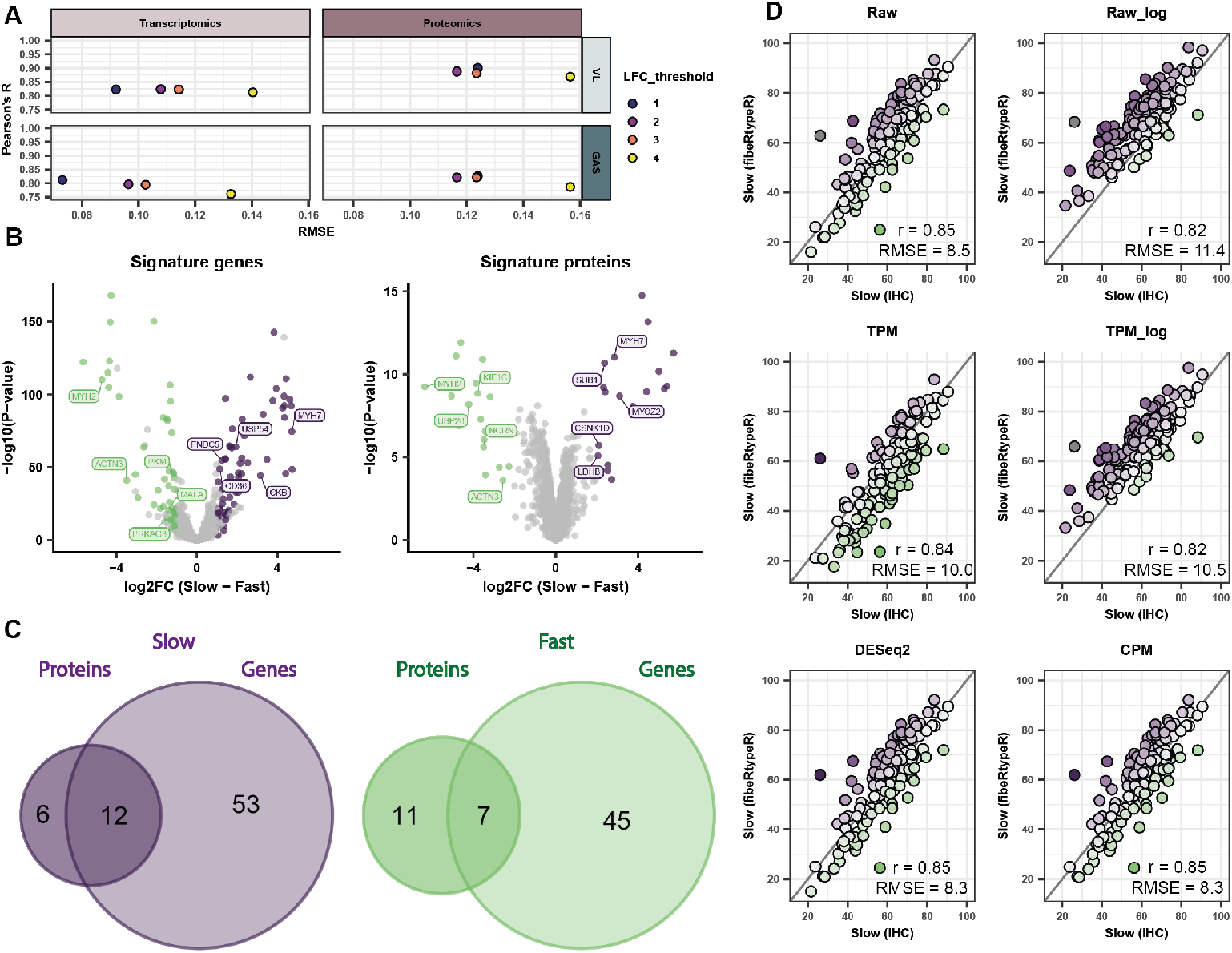
Signature matrix creation for FibeRtypeR. (**A**) Correlation coefficients (r value) and root mean square error (RMSE) for different log_2_ fold-change cut-offs in the transcriptome and proteome dataset, separately for vastus lateralis and gastrocnemius. (**B**) Volcano plots highlighting fiber type markers in transcriptomics and proteomics, based on the selected log_2_ fold changes from A. LFC cutoff of 1 for transcriptomics and 2 for proteomics. (**C**) Overlap of fiber type markers in transcriptomics and proteomics for slow and fast fibers. (**D**) Transcriptome-based FibeRtypeR accuracy with different normalization strategies for transcriptomics datasets. Diagonal lines are lines of identity.

### FibeRtypeR is methodologically robust

A key consideration in transcriptomics data analysis is the selection of data normalization strategies. We therefore benchmarked various normalization strategies in our transcriptomics workflow (no normalization, log_2_-transformation, TPM-normalization, TPM + log_2_-transformation, DESeq2, or CPM) using the previously selected list of signature genes (**Figure 3D**). TPM and both log_2_-transformation strategies resulted in the poorest correlation coefficients with IHC-based fiber type estimates, whilst DESeq2 and CPM normalization yielded the highest and nearly identical performance (r = 0.85; RMSE = 8.3). We therefore implemented CPM normalization for transcriptomics-based deconvolution in FibeRtypeR. The transcriptomics data is 3’ end sequencing data of polyadenylated transcripts with one read per transcript, irrespective of length, likely explaining why CPM works well.

Due to rapid technological developments, MS-based proteomics instruments and computational pipelines are constantly undergoing updates and modifications. Because of this, it is common that instrument combinations, acquisition methods and settings diverge between laboratories and experiments. For this reason, we evaluated if FibeRtypeR could robustly infer fiber type proportions across different MS-based proteomics workflows. A detailed comparison of acquisition methods and quantification strategies is beyond the scope of this manuscript; we further refer the reader to other detailed review articles (Cox and Mann, 2008; Sinha and Mann, 2020). We first compared FibeRtypeR results generated by data-dependent (DDA) and data-independent acquisition (DIA) modes (**Figure 4A**). Despite our reference dataset stemming from DIA-based proteomics, FibeRtypeR can successfully deconvolute DDA-based data processed using the MaxQuant software (**Figure 4B – middle panel**). However, the quantification strategy substantially affects FibeRtypeR accuracy, as using ‘unique + razor peptides’ for quantification greatly underestimated the proportion of slow fibers when compared against the values determined by IHC (r = 0.7; RMSE = 27.7; mean diff. = -25.9) (**Figure 4B – left panel**). Instead, relying only on ‘unique peptides’ for quantification markedly increased the deconvolution accuracy (r = 0.74; RMSE = 9.7; mean diff. = -0.3) (**Figure 4B – middle panel**). Not surprisingly, highest correlations were observed when applying FibeRtypeR to DIA data (r = 0.89; RMSE = 11.7; mean diff. = +9.9), either due to the reference data being DIA-based or to a larger protein coverage when comparing DIA to DDA approaches, although it also had a consistent overestimation of slow fiber proportion compared to the DDA unique peptides combination (**Figure 4B – right panel**). We next evaluated the robustness of FibeRtypeR across different MS instruments and DIA analysis software (**Figure 4C**). Overall, FibeRtypeR showed strong significant correlation coefficients when compared to the reference IHC-based fiber type proportions independently of the instrument and software of choice (r ≥ 0.85). We observed a consistent overestimation of the slow fiber fraction from timsTOF deconvolution data using unique peptides for quantification, in both DIA-NN and Spectronaut software (r = 0.85; RMSE = 11.3; mean diff. = +8.8 and r = 0.89; RMSE = 11.7; mean diff. = +9.9) (**Figure 4D – bottom middle and right panel**). This overestimation was not observed for Orbitrap Exploris data where unique peptide-based quantification produced higher accuracy in both DIA-NN (r = 0.89; RMSE = 6.6; mean diff. = -1.8) and Spectronaut (r = 0.89; RMSE = 7.1; mean diff. = +0.99) software (**Figure 4D – top middle and right panel**). Unlike with DDA data, using unique + razor peptides for quantification resulted in only a small increase of the RMSE and mean difference in the Orbitrap Exploris results (RMSE = 10.4; mean diff. = +7.4), while using unique + razor peptides for quantification actually reduced the RMSE value when using timsTOF data (RMSE = 7.8; mean diff. = +3.2) (**Figure 4D – left panels**). Despite these technical differences, FibeRtypeR consistently produced robust fiber type estimates with strong correlations to IHC across all tested MS workflows. This confirms that the method is broadly applicable to bulk MS-based proteomics datasets from human skeletal muscle biopsies, independent of platform or raw data processing pipelines.

**Figure 4.**
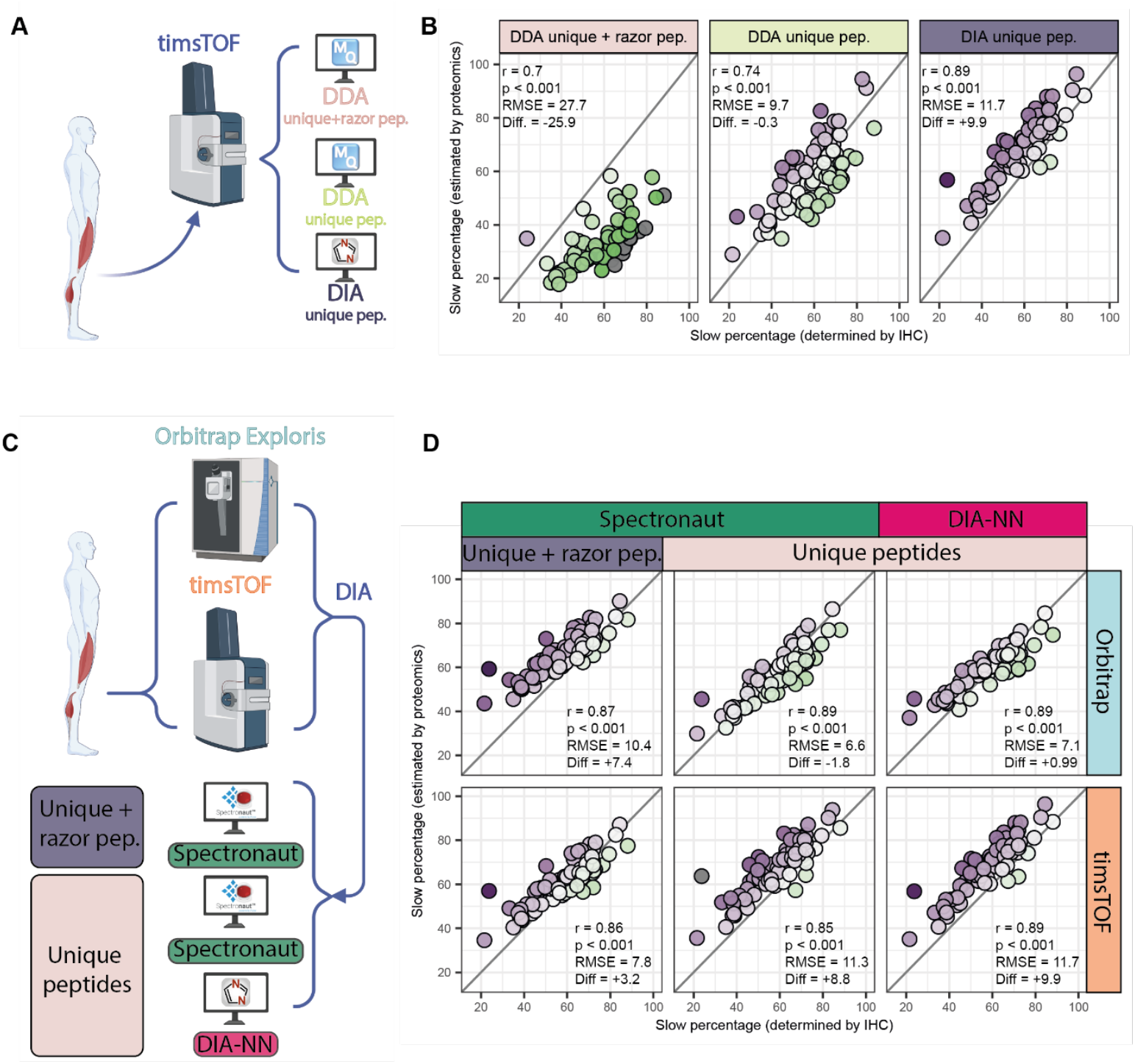
FibeRypeR is valid across proteomics acquisition and analysis pipelines. (**A-B**) Validation of FibeRtypeR with data-dependent (DDA) versus data-independent (DIA) acquisition using a timsTOF mass spectrometer. For DDA, data was quantified using either ‘unique + razor peptides’ or ‘unique peptides’ only. (**C-D**) Validation of FibeRtypeR using DIA on different instruments (Orbitrap Exploris vs timsTOF) and different software for analysis (Spectronaut vs DIA-NN). Diagonal lines are lines of identity. Diff: Average percentage-point difference compared to IHC-based fiber type based on Bland-Altman analysis.

### Applicability of FibeRtypeR

To showcase the applicability of our deconvolution tool, we applied FibeRtypeR to publicly available datasets to validate our results in biological scenarios known to affect fiber type proportions. We first tested our tool in the context of aging on a bulk *vastus lateralis* transcriptomics dataset of younger (< 50 years) and older (> 60 years) individuals (Tumasian et al., 2021). According to FibeRtypeR, older individuals had a slower typology (34.5% vs 24.1% slow fibers, p=0.002; Wilcoxon rank-sum test) compared to younger individuals (**Figure 5A**), in line with traditional fiber typing methods (Dowling et al., 2023; Miljkovic et al., 2015; Nilwik et al., 2013). Next, we investigated potential sex differences as men usually exhibit a more pronounced fast fiber type distribution than women (Haizlip et al., 2015; Nuzzo, 2024). Using a large transcriptomics dataset from 803 individuals (Melé et al., 2015), we could indeed identify a higher proportion of slow fibers in females (69.0 vs 59.7 % slow fibers, p<0.001; Wilcoxon rank-sum test) using FibeRtypeR (**Figure 5B**). We then set to investigate whether FibeRtypeR could be relevant to the study of metabolic diseases by applying our workflow to a proteomics dataset comprising skeletal muscle biopsies from normal glucose-tolerant individuals and individuals with type 2 diabetes (Kjærgaard et al., 2025). In line with previous research (Albers et al., 2015; Stuart et al., 2013), FibeRtypeR identified a significant difference in the fiber type proportion between groups, with individuals with type 2 diabetes displaying a lower abundance of slow fibers (50.4% vs 55.7% slow fibers, p=0.002; unpaired Welch’s T-test) (**Figure 5C**). Moreover, the fiber type proportions extracted from FibeRtypeR were significantly associated with insulin sensitivity (hyperinsulinaemic-euglycaemic clamp-derived M-value; r = 0.413, p < 0.001), highlighting its potential in analyzing clinically relevant phenotypes (**Figure 5D**).

**Figure 5.**
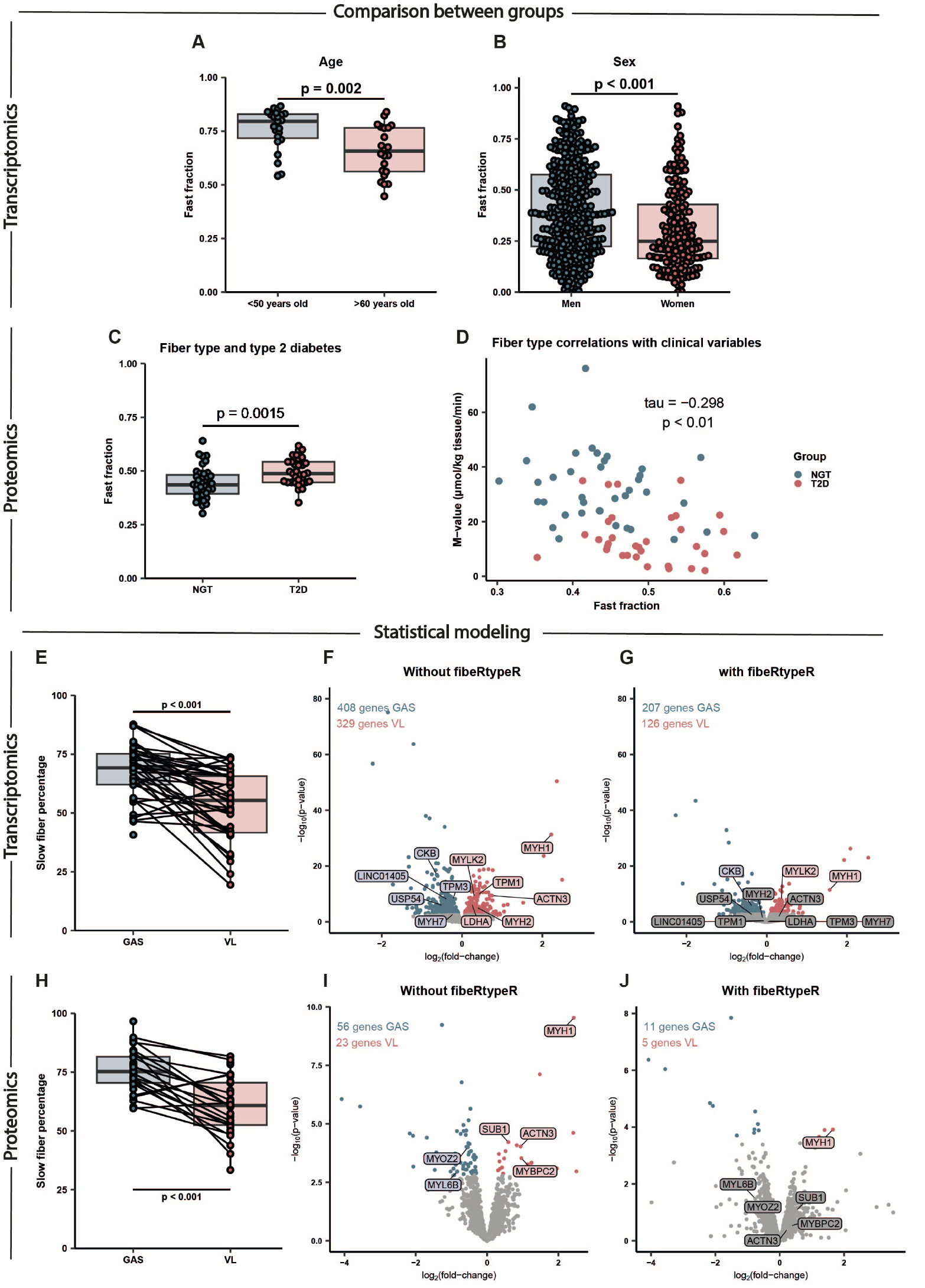
Application potential of FibeRtypeR. (**A-B**) Transcriptome-based fiber type inference in (A) young (<50 years) and old (>60 years) or (B) male and female vastus lateralis samples. (**C-D**) Proteome-based fiber type inference from vastus lateralis samples in individuals with normal glucose tolerance (NGT) and type 2 diabetes patients (T2D), and its correlation with the individual M-value (reflective of insulin sensitivity). (**E-G**) Transcriptome-and proteome-based fiber type inference of gastrocnemius (GAS) and vastus lateralis (VL) samples. (**G-H**) Differentially expressed/abundant genes/proteins in GAS and VL samples without using fiber type proportions as a covariate in the statistical pipeline. (**I-J**) Differentially expressed/abundant genes/proteins in GAS and VL samples with using fiber type proportions as a covariate in the statistical pipeline.

Alternatively, FibeRtypeR can be valuable to account for variation in fiber type proportions in statistical modelling, such as in differential expression analysis. This is especially relevant since human muscle biopsy sampling is characterized by substantial inter-biopsy variability, which is usually not taken into account (Horwath et al., 2021; Van de Casteele et al., 2024b). As proof-of-principle, we divided the transcriptome and proteome datasets into two groups based on which muscle was sampled, *vastus lateralis* or *gastrocnemius*. As expected, FibeRtypeR detected a higher slow fiber proportion in gastrocnemius muscle for both transcriptomics (**Figure 5E**) and proteomics (**Figure 5F**). Importantly, failing to account for fiber type composition in differential gene and protein expression analysis led to numerous features being differentially expressed between muscles, many of which (e.g., MYH2, MYH7 and MYL6B) are directly linked to fiber type (**Figure 5G-H**). Simply accounting for fiber type proportions obtained from FibeRtypeR in the statistical model drastically reduces the number of differential features, allowing for assessment of inherent differences between muscles independent of fiber type composition (**Figure 5 I-J**).

### FibeRtypeR is available as an easy and open-source Shiny app

To make fiber type deconvolution accessible to the broader skeletal muscle research community, we developed FibeRtypeR as a free and user-friendly Shiny web application available at https://muscleapps.ugent.be. The app seamlessly integrates the deconvolution workflow with dedicated tabs for instructions on data formatting, data upload and fiber type inference (**Figure 6A**). Users are provided with clear guidance on how to structure their input data, including a downloadable template file to assist with formatting (**Figure 6B**). The app accepts raw count data for transcriptomics and raw LFQ intensity values for proteomics, which are then real-time normalized upon selection of the data type (**Figure 6C**). The actual deconvolution is then initiated via a single-click of a button (“Run FibeRtypeR”). The algorithm executes rapidly (1-10 seconds), and results are immediately displayed as a figure and a table, which can be downloaded (.png or.html files for the figure;.csv,.tsv or.xlsx for the table) for further analyses by the user (**Figure 6D**).

**Figure 6.**
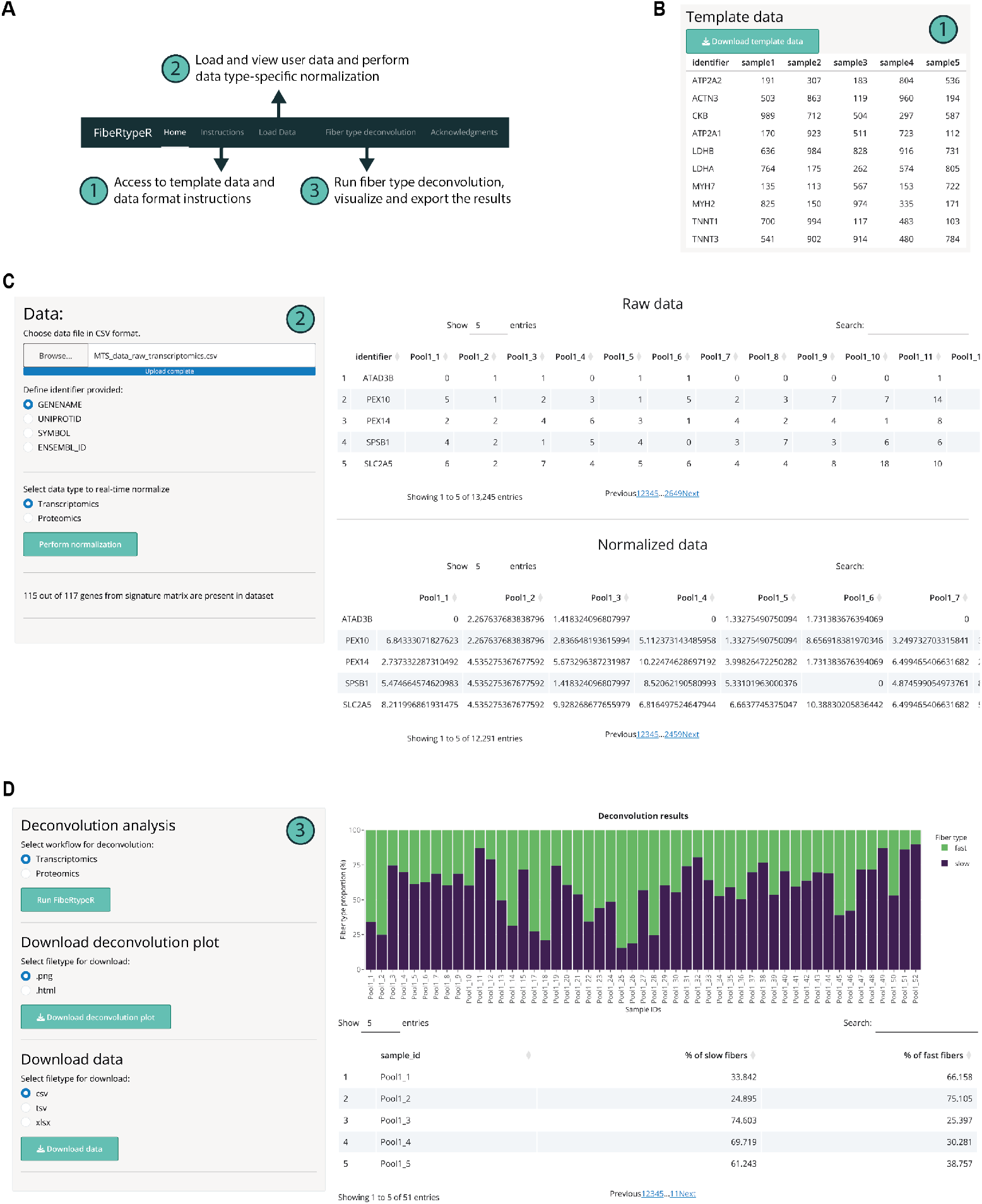
FibeRtypeR is available as an open-access Shiny app. (**A**) Pipeline of FibeRtypeR with (1) clear instructions for data formatting, (2) quick data-specific normalization and (3) options for data visualization and download. (**B**) Available template data to help with data formatting. (**C**) Raw data is uploaded and is then automatically normalized for further use in FibeRtypeR. (**D**) FibeRtypeR gives accurate results within seconds, including visualization of results, and allows for data download by the user.

## Discussion

Here we developed FibeRtypeR, an easy-to-use open-source online tool to estimate fiber type proportions from bulk transcriptomics and proteomics datasets. We leveraged our previously generated omics profiles of approximately 1000 individual human skeletal muscle fibers for both transcriptomics and proteomics to allow an accurate fiber type inference from independent bulk datasets (Moreno-Justicia et al., 2025). We validated FibeRtypeR by comparing it to paired data of IHC-determined fiber type (only MYH-based), and applied FibeRtypeR to previously published datasets, resulting in the identification of fiber type shifts with aging and type 2 diabetes, or fiber type differences between sexes and muscle groups.

While both the transcriptome and proteome workflow yielded strong agreement with the IHC-based fiber type proportions, FibeRtypeR estimated a greater proportion of slow fibers using proteomics data than what would be determined via the immunohistochemical analysis of MYH isoforms alone. This overestimation of the slow fiber type fraction only happened when using proteomics data derived from measurements in a timsTOF instrument or using ‘unique and razor peptides’ for quantification in an Orbitrap instrument. These differences suggest that the mass analyzer and the presence or absence of ion mobility within the instrument may play a significant role in quantifying protein isoforms important for fiber type determination. Nonetheless, the observed differences could be in part related to a substantial slow MYH7 protein expression in most fast fibers and discordances in RNA expression and protein abundance, as recently exemplified in the absence of pure type 2X fibers in the proteome (Bishop et al., 2023; Moreno-Justicia et al., 2025; Vogel and Marcotte, 2012). These results therefore raise important questions as to whether fiber-typing using a single classifier can be considered the true gold standard, especially when employed via IHC, which is only semi-quantitative and less specific than RNA sequencing or mass spectrometry. In contrast, FibeRtypeR takes into account 117 genes and 36 proteins to determine fiber type proportions. Well-described contractile genes and proteins with fiber type specific expression (e.g., MYH2/MYH7, TNNI2/TNNI1 and TNNC2/TNNC1) are included in the signature matrix on which the deconvolution algorithm is based (Murgia et al., 2021; Rubenstein et al., 2020). However, features related to other biological pathways are also integrated, including glucose metabolism, ubiquitin regulation and non-coding RNAs. FibeRtypeR therefore likely captures a more complete spectrum of phenotypic and functional differences between fiber types. Notwithstanding the above limitations of IHC-based fiber typing, the degree of bias between FibeRtypeR and the current gold standard approach (IHC) is modest (0-9 percentage points) and consistent, allowing for the effective application of FibeRtypeR to identify differences in fiber type proportions between groups, such as with aging and between biological sexes. FibeRtypeR is also capable of identifying muscle-specific differences in fiber type proportions, as the *gastrocnemius* exhibited a greater proportion of slow fibers compared to the *vastus lateralis* (Houmard et al., 2000), and this was consistent across the three fiber typing methods.

Deconvoluting fiber type proportions from bulk omics datasets is not entirely novel (Oskolkov et al., 2022), although our approach presents several improvements and expansions. Firstly, the FibeRtypeR reference datasets are built from whole fiber transcriptomics and proteomics profiles, whereas this earlier endeavour was based on single-nuclei RNA-seq data, characterized by only shallow sequencing and not taking into account the multinucleated nature of skeletal muscle fibers. Secondly, our reference dataset is representative of multiple individuals whereas this previous approach relied upon data from a single individual. Furthermore, our approach is also compatible with mass spectrometry-based proteomics as is often used in muscle physiology and exercise research (Cervone et al., 2024). Finally, we provide a point-and-click interface for users to determine fiber type proportions from their own datasets within seconds, which will be a valuable resource for the muscle research community.

In conclusion, we present FibeRtypeR, a validated and time-efficient deconvolution tool to accurately estimate fiber type proportions from bulk skeletal muscle transcriptome and proteome datasets. FibeRtypeR is built upon a robust reference dataset of approximately 1000 fibers from multiple donors for both transcriptomics and proteomics applications. FibeRtypeR provides an effective tool to go beyond myosin heavy chains in the analysis of fiber types in the era of omics research.

## Methods

### Bulk muscle validation dataset

#### Participants

Forty healthy (20 female), young participants underwent four biopsies in total. They were involved in various recreational or competitive sports for at least 2 h/week. Subjects were instructed to refrain from alcohol, caffeine and exercise in the 24 h preceding the biopsy sampling. This study was conducted according to the Declaration of Helsinki and was approved by the local ethics committee of Ghent University Hospital, Belgium (Ethics Application BC-09442). All participants gave written informed consent to partake in this study.

#### Muscle biopsy collection

Participants reported to the lab on one occasion and consecutively underwent two biopsies in the *vastus lateralis* and two in the *gastrocnemius medialis*. Muscle biopsy samples were collected using a 5 mm Bergström needle with suction (Bergstrom, 1975, 1962) under local anaesthesia (0.5 ml xylocaine, Aspen Netherlands, Gorinchem, The Netherlands), as described previously (Van de Casteele et al., 2024b). Immediately after, the muscle fiber bundles of the sample(s) were oriented perpendicularly, mounted in embedding medium (OCT Compound, Tissue-Tek, Sakura Finetek, Zoeterwoude, The Netherlands), frozen in liquid nitrogen-cooled isopentane and eventually frozen in liquid nitrogen. All muscle samples were stored at −80 °C until processing.

#### Immunohistochemistry and imaging

All embedded muscle samples were cut into 8 μm muscle cross-sections using a cryostat (Epredia HM560, Machelen, Belgium). Thereafter, the embedded muscle samples were mounted on microscopy glass slides and stained for the muscle fiber membrane (wheat germ agglutinin), type 1 fibers (BA-F8), type 2A fibers (A4.74) and type 2X (6H-1) fibers. Slides were then imaged using a fluorescence microscope with a ×10 objective (Zeiss Axioscan 7, Zeiss, Zaventem, Belgium), and fiber type proportions and cross-sectional area (fCSA) were measured using ImageJ software (National Institutes of Health, Bethesda, MD, USA) with Fiji extension. Type 2A, 2A/X and 2X fibers were all classified as fast fibers, and type 1/2A (< 1%) were excluded for further analysis. The relative area occupied by type 1 fibers was used to validate FibeRtypeR and was calculated by multiplying the average type 1 fCSA by the number of type 1 fibers divided by the total cross-sectional area of type 1 and type 2 fibers. A detailed description of our immunohistochemical methodology has been provided previously (Van de Casteele et al., 2024b).

#### RNA sequencing

All 160 samples were used for bulk RNA sequencing. Most samples (73%) were snap-frozen tissue, while the other samples (27%) were initially embedded in OCT compound (Tissue-Tek, Sakura Finetek, Zoeterwoude, The Netherlands), which was manually removed before RNA isolation. Total RNA was then isolated from the muscle biopsies using the miRNeasy Mini kit (Qiagen, #217004) according to the manufacturer’s instructions, with 30 µL elution volume. RNA integrity was checked (FragmentAnalyzer), followed by sample dilution to equal concentrations after NanoDrop quantification and gDNA removal with 60 ng RNA as input in a 12 µL reaction (HL-dsDNase kit, #70800, ArcticZymes). Illumina-compatible libraries from 2 µL DNase-treated RNA were then prepared using the QuantSeq-Pool 3’ mRNA-Seq library prep kit (Lexogen), with 12 cycles for endpoint PCR. Quality of libraries were checked (high sensitivity small DNA Fragment Analysis kit, Agilent Technologies, DNF-477-0500), followed by equimolar pooling (4 nM) and sequencing with a NovaSeq S1 kit (v1.5, #20028319) on a NovaSeq 6000 instrument with loading of 1.90 nM (2% Phix). Primary data analysis was performed with the QuantSeq-Pool analysis pipeline (https://github.com/Lexogen-Tools/quantseqpool_analysis). Genes with fewer than 50 total raw counts across samples were removed and one sample was omitted during quality control (distance and principal component plots).

#### MS-based proteomics

Only a subset of the samples used for RNA sequencing were analyzed using MS-based proteomics (n = 77). Muscle lysates were prepared by powdering of snap-frozen (n = 46) and initially embedded (n = 31) muscle biopsies before resuspending in lysis buffer (1% sodium deoxycholate, 50 mM Tris PH 8.5), followed by homogenization using a tissue homogenizer (IKA turrax). Next, lysates underwent boiling at 95 °C in a thermomixer (Eppendorf) for 10 min followed by sonication using a tip sonicator. After protein concentration determination using BCA (Thermo Fisher), 40 μg of protein were alkylated and reduced upon addition of DTT and CAA to final concentration of 10 mM and 40 mM, respectively. Proteins were digested overnight by addition of LysC and trypsin (1:500 and 1:100 enzyme to protein ratio, respectively) in a thermomixer at 36 °C. The next morning, tryptic peptides were desalted using in-house made SDB-RPS stage-tip columns prior loading of 200 nanograms of peptides into Evotips (Evosep One) according to the manufacturer’s instructions. Samples were run in data-independent or data-dependent parallel accumulation serial fragmentation (DIA-PASEF or DDA-PASEF) mode on an Evosep One LC-system (Evosep, Denmark), in-line connected to a timsTOF SCP (Bruker). Peptides were analyzed with the 60 SPD method using the EV-1064 endurance column (8 cm x 100 µm I.D., 3 µm beads, Evosep, Denmark), heated to 45 °C. Peptides were eluted from the column through the predefined 60SPD gradient consisting of 0.1% FA in LC-MS-grade water as solvent A and 0.1% FA in ACN as solvent B. For DIA-PASEF, eluting peptides were measured in positive polarity with a full-scan range of 100 m/z to 1700 m/z. The TIMS was operated at a fixed duty cycle close to 100%, a ramp and accumulation time of 100 ms, ranging from 1/K0 = 0.64 Vscm2 to 1/K0 = 1.50 Vscm2. Collision energy was linearly ramped as a function of the inverse mobility from 20 eV at 1/K0 = 0.60 Vscm^2^ to 59 eV at 1/K0 = 1.60 Vscm^2^. A DIA-PASEF mass range of 400 Da to 1000 Da was used in a mobility range of 1/K0 = 0.64 Vscm^2^ to 1/K0 = 1.37 Vscm^2^, resulting in a cycle time of 0.96s. For DDA-PASEF, a ten PASEF/MSMS scan acquisition method was used in DDA-PASEF mode with a precursor signal intensity threshold at 500 arbitrary units. An adapted polygon in the m/z-IM plane was used (IMS filter mass: IMS filter mobility; 270.54: 0.551; 400.74: 0.850; 702.59: 1.101; 704.02: 1.723; 1716: 1.764). The mass spectrometer was operated in non-sensitive mode with an accumulation and ramp time of 100ms, analyzing in MS from 100 to 1700 m/z. Precursors were isolated with a 2 Th window below m/z 700 and 3 Th above and actively excluded for 0.4 min when reaching a target intensity threshold of 20,000 arbitrary units. A range from 100 to 1700 m/z was covered with the following collision energy scheme (Vs cm^-2^ / eV; 0.7 / 20; 1.06; 30; 1.16; 40; 1.34; 40; 1.68; 70).

From here, raw MS files were processed using the DIA-NN (*v. 1.8.1*) (Demichev et al., 2022) or Spectronaut (*v. 18*) (Biognosis) software for DIA runs, while the DDA runs were underwent processing via the MaxQuant (*v. 2.6.*4) pipeline. DIA-NN searches were conducted using proteotypic peptides for protein group quantification and double-pass mode was chosen as neural network mode, “Robust LC (high accuracy)” was set as quantification strategy and the match between runs options was enabled, the rest of the parameters remained as default, including precursor FDR set to 1% and peptide length of 7-30 amino acids. Spectronaut searches were conducted twice, using the default parameters for quantification first and then swapping the quantification mode to use only unique peptides for quantification. The rest of the software specification remained as the default of the directDIA + mode. Similarly, two MaxQuant DDA searches were run in parallel with different quantification settings. Under Global parameters, in the protein quantification tab, one search used “Unique + razor” while “Unique” was selected for the other search. The remaining parameters were left as default, after specifying “Bruker tims” as an instrument.

#### Single-fiber transcriptomics and proteomics reference datasets

Transcriptome and proteome profiles of individual human skeletal muscle fibers were used from our previous analysis (Moreno-Justicia et al., 2025). For transcriptomics, fibers were isolated from fourteen young and healthy individuals (n = 925 fibers), while 974 fibers from 5 different individuals were collected for the proteomics dataset. A slow or fast type was then assigned to each fiber based on the UMAP clustering, resulting in two distinct clusters (i.e., slow and fast fibers). Subject-specific pseudobulk profiles were then generated by aggregating counts (transcriptomics) or taking the median of LFQ intensities (proteomics) of each slow and fast fiber separately. This pseudobulked data was used for further processing as described below.

#### Transcriptomics application datasets

A skeletal muscle transcriptomics dataset of 53 healthy individuals with an age range of 22-83 years old was downloaded from Gene Expression Omnibus with accession number GSE164471 (Tumasian et al., 2021). Genes with fewer than 10 median raw counts across samples were filtered out and remaining raw data were normalized using the ‘counts per million’ strategy. Subjects were then classified in a young (<50 years, n = 23) and old (> 60 years, n = 22) group. Estimated fiber type proportions were compared between groups with a Wilcoxon rank-sum test. For the analysis of sex differences, a large skeletal muscle dataset of 818 individuals (20-80 years old) was downloaded from the GTEx Portal (Melé et al., 2015). Raw counts were normalized with ‘counts per million’ after excluding genes that had less than 8030 total raw counts across all samples. Fiber type proportions were compared between men (n = 552) and women (n = 266) with a Wilcoxon rank-sum test test.

#### Proteomics application dataset

Raw LFQ intensities from *vastus lateralis* of NGT (n = 39) and T2D (n = 33) individuals, as well as anonymized sample information were retrieved (Kjærgaard et al., 2025). After loading the data in FibeRtypeR, 21 protein markers out of 36 were present in the data and used for fiber type deconvolution. The resulting fiber type proportions were loaded in RStudio and compared between groups using an unpaired Welch’s T-test. Fast fiber type proportions were correlated against the individual’s M-value using Kendall’s tau.

### Deconvolution analysis

#### Signature matrix

Signature matrices specific for each omics technology were generated based on pseudobulked differential expression results comparing slow and fast fibers (Moreno-Justicia et al., 2025). Genes and proteins with log_2_ fold changes greater than one (transcriptomics) and two (proteomics) were selected as excellent markers for both fiber types and were used as reference input in the deconvolution pipeline. This resulted in 65 and 52 genes for slow and fast fibers and 18 and 18 proteins for slow and fast fibers. Genes that are responsive to acute exercise were also omitted from the signature matrix. To this end, we reanalyzed our previously published bulk RNA-seq dataset (Van der Stede et al., 2025), and labelled a gene as exercise-responsive if its absolute log_2_ fold change was greater than 0.58 (fold change of ±1.5) at either immediately post-exercise or after 3 hours of recovery. Expression of these genes were then averaged for slow and fast fibers across participants, based on the cpm-normalized and pseudobulked transcriptomics dataset, as well as for the log_2_-normalized LFQ intensities in the proteomics dataset. This resulted in two values per gene/protein, being the representative expression of those genes/proteins in slow and fast fibers, respectively. Overlap of genes/proteins per fiber type were explored with the *eulerr* package (v. 7.0.0). For selecting the best transcriptomics normalization strategy, results were compared using no normalization (*raw*), log_2_-transformed counts (*raw_log*), TPM normalization (*TPM*), log_2_-transformed TPM counts (*TPM_log*), DESeq2-based normalization (*DESeq2*) and ‘counts per million’ normalization (*cpm*).

#### Deconvolution

The deconvolution analysis was performed with the *DeconRNASeq* package (v. 1.42.0) in RStudio, with as input data both the normalized bulk and fiber type specific reference data (checksig = F, known.prop = F, use.scale = F). Correlations between histochemical determination of fiber type (% cross-sectional area) and omics-based inference were done with Pearson correlations with the *cor.test()* function. Bland-Altman plots were constructed for *vastus lateralis* and *gastrocnemius medialis* to visualize and evaluate the bias for mean differences between fiber type determined by IHC and estimated by transcriptomics or proteomics. The agreement interval was calculated by adding and subtracting 1.95*standard deviation to the mean difference. Root mean square errors were determined with the *rmse()* function from the *Metrics* package (v. 0.1.4). Differential gene expression analysis between *vastus lateralis* and *gastrocnemius* muscle was performed with *DESeq2* (v. 1.40.2) with either a “∼ group” (without FibeRtypeR) or “∼ fiber type + group” (with FibeRtypeR) statistical model. In turn, the *limma* package (v. 3.58.1) was used for the differential protein abundance analysis between *vastus lateralis* and *gastrocnemius* muscles, with the following statistical model: “∼ 0 + muscle + subject” in the without FibeRtypeR comparison, and adding fast fiber type percentage as a covariate in the with FibeRtypeR comparison.

## Data availability

All RNA sequencing and proteomics data will be made publicly available upon acceptance of the manuscript.

## Acknowledgments

Bulk muscle mass spectrometry analyses were performed at the VIB Proteomics Core (Ghent, Belgium). This work was generously funded by grants from the Research Foundation Flanders (G080321N, 11B4220N, 1S66923N; to WD, TVdS and FVdC, respectively). The Novo Nordisk Foundation Center for Basic Metabolic Research is an independent Research Center, based at the University of Copenhagen, Denmark, and is partially funded by an unconditional donation from the Novo Nordisk Foundation (www.cbmr.ku.dk; grant numbers NNF18CC0034900 and NNF23SA0084103). Several illustrations were made with BioRender.com. The technical assistance of Ruud Van Thienen, Anneleen Weyns, Anneke Volkaert, Nurten Yigit, and Jasper Anckaert is greatly appreciated.

## Author contributions

Conceptualization: TVdS, RMJ, FVdC, BS, ASD, WD

Methodology: TVdS, RMJ, FVdC, MH, PM, JVS, BS, ASD, WD

Software: TVdS, RMJ, PMer

Validation: TVdS, RMJ, FVdC, JVS, ASD, WD

Formal Analysis: TVdS, RMJ, FVdC

Investigation: TVdS, RMJ, FVdC, EL, AVdL, JVdc, DVH, AS, BS

Resources: JVdE, SD, MH, PM, JVS, ASD, WD

Data curation: TVdS, RMJ, FVdC, BS

Writing – Original Draft: TVdS, RMJ, FVdC, BS, ASD, WD

Writing – Review and Editing: TVdS, RMJ, FVdC, EL, AVdL, JVdc, PMer, JVdE, DVH, AS, SD, MH, PM, JVS, BS, ASD, WD

Visualization: TVdS, RMJ, FVdC

Supervision: SD, PM, JVS, BS, ASD, WD

Project Administration: TVdS, RMJ, FVdC, BS, JVS, ASD, WD

Funding Acquisition: TVdS, FVdC, JVdE, PM, JVS, ASD, WD

